# Investigating the relationship Between AMBRA1 and Cell Proliferation in Melanoma

**DOI:** 10.1101/2025.07.29.667464

**Authors:** Milad Ibrahim, Marco Corazzari, Jane Armstrong, Noel Carter

## Abstract

The protein Activating molecule in Beclin1-regulated autophagy1 (AMBRA1), discovered in 2007, is crucial for autophagy and plays roles in nervous system development, cell survival, and proliferation. This study investigates AMBRA1’s involvement in various cellular processes using a systems-based “omics” approach, focusing on melanoma. Transcriptomic analysis of the overexpression or the knock-down of AMBRA1 was shown to result in significant dysregulation of several transcripts. This appears to have identified several novel roles for AMBRA1 in a range of cellular pathways; some of these are hallmarks of cancer signaling including, MAPK, angiogenesis, tissue growth factor signaling, axon guidance and Wnt signaling. Yeast two-hybrid assays performed in this study identified novel binding partners that can provide evidence for new roles for AMBRA1 in different cellular processes. The work shows that AMBRA1 loss appears to upregulate metastatic genes/proteins supporting the role of AMBRA1 as a tumor suppressor gene in melanoma.

## Introduction

Melanoma accounts for 75% of skin cancer deaths and remains one of the most therapy-resistant and aggressive cancers. Survival rates for five years after being diagnosed with melanoma are highly dependent on the stage of the disease. If not diagnosed and treated early, melanoma can become highly metastatic. Globally, cases of melanoma are predicted to rise by approximately 50% by 2040 [1, 2]. Considerable improvement in melanoma treatment has been achieved using immunotherapy, targeted therapies, and combination therapies. Despite advancements in melanoma treatment, metastatic disease response remains around 50% [3], highlighting the need for personalized therapy based on the molecular features of tumors remains crucial to achieving better clinical outcomes [4, 5].

Activating Molecule in BECN1-regulated autophagy 1 (AMBRA1) is a scaffold protein that plays a central role in regulating a range of cellular processes. Its primary identified function is the regulation of autophagy and the development of the nervous system [6]. Advances in the understanding of this protein have shown that it is involved in key physiological events such as metabolism, apoptosis, and cell proliferation. It is also an important regulator of embryonic development [7]. *AMBRA1* has been shown to have a tumor suppressor role in some cancers and a pro-tumorigenic role in others. For example, AMBRA1 loss in the tumor microenvironment is associated with melanoma metastasis [8, 9] and its loss in the tumor, can promote melanoma invasion in mouse models [10]. The role of *AMBRA1* as a tumor suppressor is supported by its role in associating with PP2CA to dephosphorylate and degrade c-MYC leading to decreased cell proliferation [11] and, its role in facilitating the degradation of Cyclin D, therefore preventing the overactivation of CDK4/6 which leads to increased cell proliferation [12, 13]. Conversely, studies suggest that higher levels of AMBRA1 have been associated with increased resistance of breast cancer to epirubicin [14], as well as tumorigenesis in breast [15] and gastric cancers [16]. It is also reported to desensitize human prostate cancer cells to cisplatin [17].

Despite recent advances in understanding the role of *AMBRA1* in cell proliferation and melanoma, it remains unknown if *AMBRA1* loss promotes tumorigenesis in melanoma by additional mechanisms. In this study, we analyzed publicly available human melanoma tumor samples microarrays, and RNA-Seq datasets to assess *AMBRA1* mutation and the levels of its expression in primary and metastatic melanoma. We also performed transcriptional analysis on *AMBRA1* overexpression and knockdown in A375 melanoma cell lines. Finally, we identified putative AMBRA1 protein binding partners. This study proposes novel roles for *AMBRA1* in a range of cellular processes suggestive of a more complex biology.

## Results

### AMBRA1 expression in melanoma

We performed a bioinformatic analysis on publicly available datasets to investigate the levels of AMBRA1 in melanoma. First, we compared AMBRA1 expression levels in primary, adjacent skin and metastatic melanoma. AMBRA1 expression was significantly decreased in adjacent skin compared to primary (*p* < 0.001) and metastatic melanoma (*p* < 0.001) as previously reported [9] (**Fig. 1A**). Next, we tested AMBRA1 expression levels and found no difference between primary and metastatic melanoma in four independent datasets (**Fig. 1A-D**). AMBRA1 expression levels did not correlate with overall survival (**Fig. 1E**) or disease specific survival (**Fig. 1F**) in the Cancer Genome Atlas Program (TCGA) Skin Cutaneous Melanoma (SKCM). *AMBRA1* mutations were reported in 10% of SKCM and melanoma is the second type of cancer that was enriched for *AMBRA1* mutations after Uterine Corpus Endometrial Carcinoma (**Fig. 1G**). *AMBRA1* mRNA levels were significantly less (*p* = 0.0093) in *AMBRA1* mutated SKCM (Fig. 1H). Together, these results indicate that a subset of melanoma tumors favor AMBRA1 loss.

**Fig. 1.**
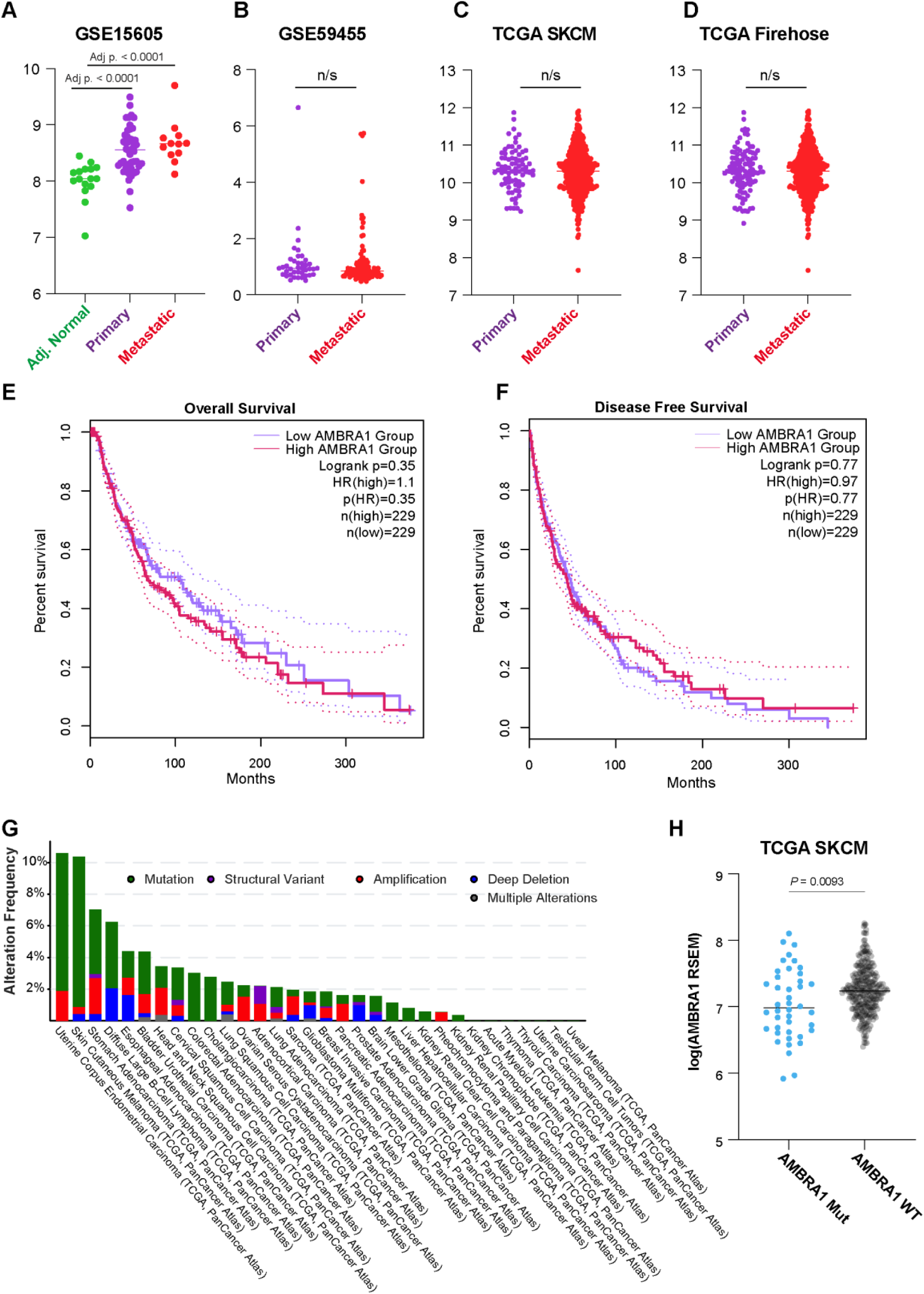
AMBRA1 expression and mutation in melanoma. (**A-D**) Dot Plot showing the expression of AMBRA1 in adjacent normal skin, primary and metastatic melanoma in 4 different melanoma datasets (A) GSE15605 (B) GSE59455 (C) TCGA SKCM (D) TCGA Firehose. *P*-values are calculated by a two-sided t-test. (**E, F**) Kaplan Meier graphs showing overall (E) and progression free (F) survival in high and low AMBRA1 expression in melanoma. *P*-values are calculated by log-rank test. (**G**) Bar plot showing AMBRA1 mutation percentage in all cancer types. (**H**) Dot Plot showing the expression of AMBRA1 in AMBRA1 mutant or wild type melanoma patients in TCGA SKCM. *P*-values are calculated by a two-sided t-test.

### Effect of AMBRA1 knockdown on melanoma cell growth

To test the effect of *AMBRA1* expression levels on cell growth, we utilized A375 melanoma cells overexpressing AMBRA1 (rAMBRA1) or A375 AMBRA1 depleted cells (shAMBRA1) (**Fig. 2A, B**). Live cell imaging assays showed that AMBRA1 overexpression did not affect growth of the melanoma cell line (**Fig 2C**). Conversely, AMBRA1 depletion led to significant decrease in melanoma cell growth over time (**Fig. 2D**). Of note, after a prolonged time of removing the shAMBRA1 construct selection antibiotic, the cells restored their ability to grow indicating that the cells favor AMBRA1 expression (**Fig. 2E**). To validate our findings, we navigated the cancer dependency map (Depmap) portal which defines which genes are essential for cancer cell survival. A negative score indicates that a gene is essential and its loss affects the proliferation of cancer cells while a positive score indicates that the gene loss does not affect cancer cell proliferation. Depmap portal showed a similar effect of *AMBRA1* knockdown using RNAi in A375 cells (**Fig. 2F)** and, knockout using CRISPR (**Fig. 2G**). Of note, Depmap RNAi and CRISPR databases showed that *AMBRA1* can be an essential gene in some melanoma cell lines. However, its loss can have no effect on other melanoma cell lines. Collectively these results suggest that the role of *AMBRA1* in melanoma cell proliferation can be cell and context dependent.

**Fig. 2.**
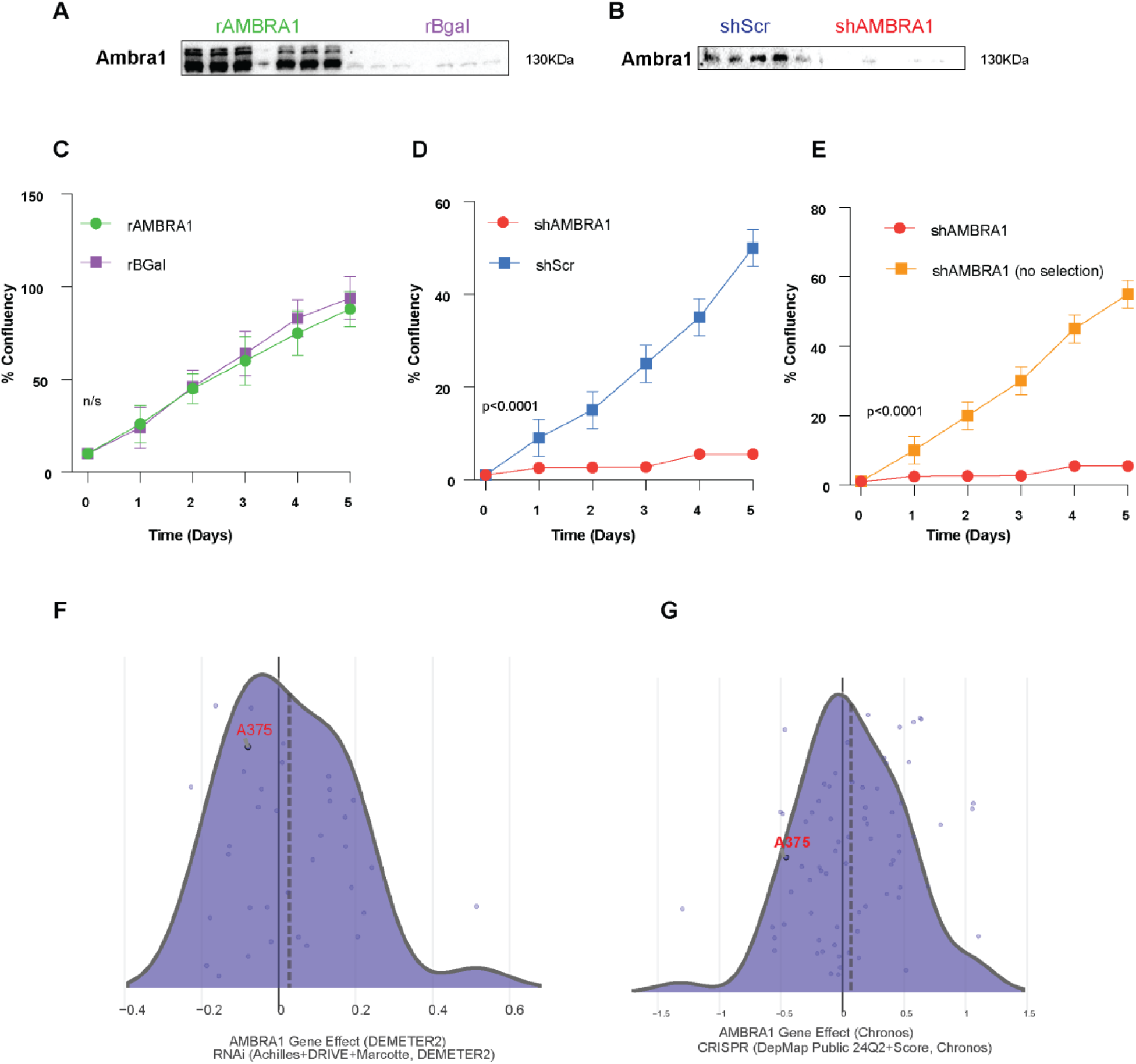
AMBRA1 loss decrease A375 melanoma cell proliferation. (**A, B**) Western blot analysis showing (A) AMBRA1 overexpression or (B) Knockdown in A375 melanoma cell lines. (**C-E)** Growth curves of A375 cell lines expressing (C) rAMBRA1 and rBgal, (D) sh AMBRA1 and shScramble or (E) shAMBBRA1 and shAMBRA1 with no selection. Data points indicate mean ± SD (n=9). *P*-values are calculated using a two-tailed t-test comparing the non-linear fit of exponential growth. **(F, G)** DepMap Chronos score plot showing the effect of AMBRA1 (F) Knockdown by RNAi or (G) knockout by CRISPR on melanoma A375 cell lines. solid line represents zero and dotted line represents median of score across cell lines.

### AMBRA1 plays a role in regulating hallmark melanoma pathways

To investigate the molecular pathways that can be regulated by *AMBRA1*, we performed transcriptomic analysis using A375 cells with either the overexpression or knockdown of AMBRA1. 3D principal component analysis (PCA) revealed that shAMBRA1 clustered separately from the other cell lines (**Fig. S1**). AMBRA1 overexpression resulted in the upregulation of 60 and downregulation of 47 transcripts (*p* <0.05, –2> log2fc >2), however, when filtering for differentially expressed transcripts with significant FDR, only AMBRA1, NAA11 and CDH13 were upregulated in cells overexpressing AMBRA1 compared to the control (**Fig 3A).**

**Fig. 3.**
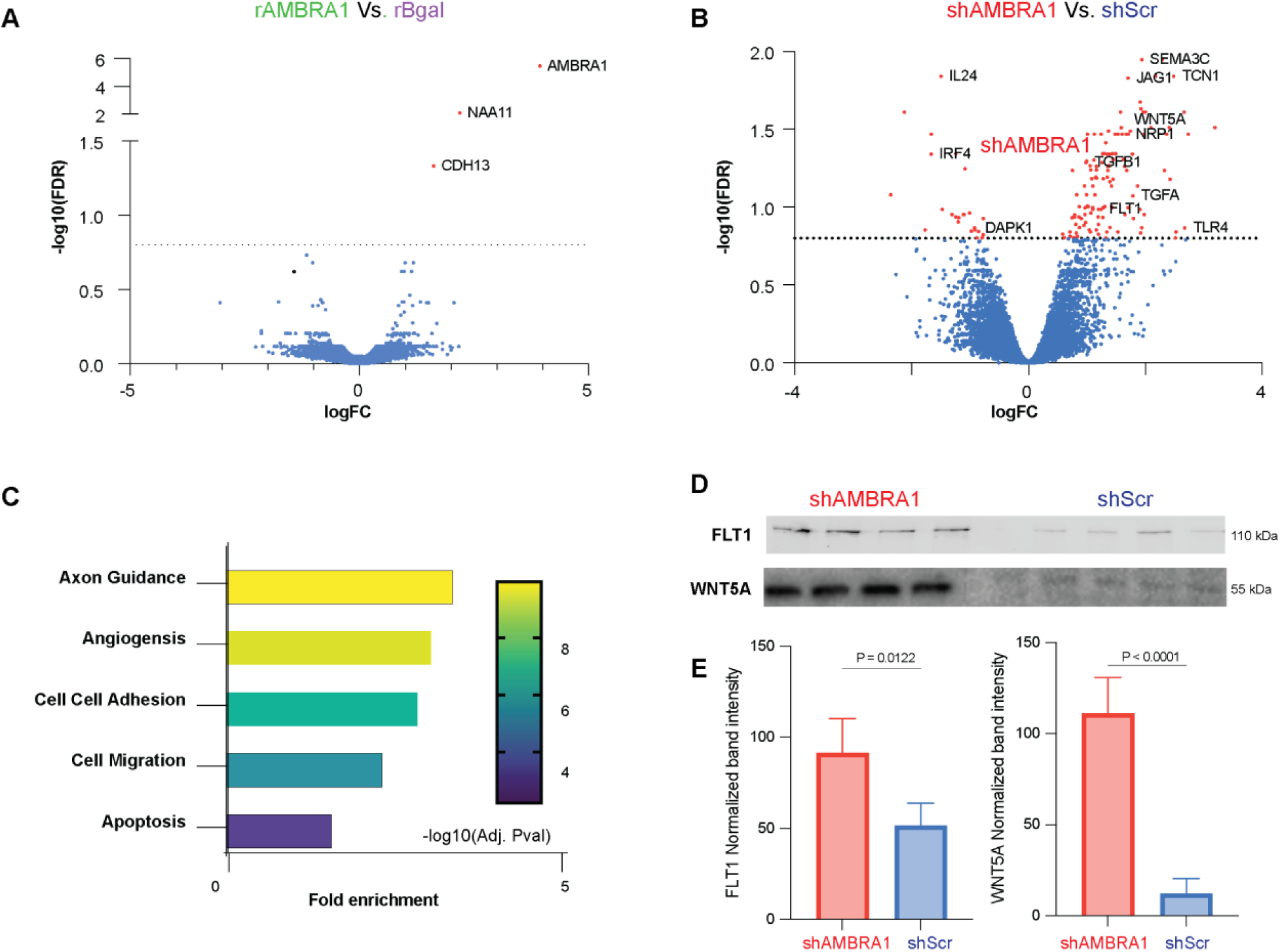
AMBRA1 regulate hallmark cellular processes on the transcriptional level. (**A**) Volcano plot showing differentially expressed genes between rAMBRA1 and rBgal. (**B**) Volcano plot showing differentially expressed genes between shAMBRA1 and shScramble (**C**) GO biological processes analysis of differential expressed genes in shAMBRA1 compared to shScramble. (**D**) Western blot analysis showing FLT1 and Wnt5A expression in A375 melanoma cell lines comparing shAMBRA1 and shScramble. *P*-values were calculated using a two-tailed t-test.

AMBRA1 knockdown resulted in 103 and 11 genes being significantly up and down regulated (p < 0.05, FDR < 0.09, –2> log2fc >2), respectively. AMBRA1 (log2FC = –1.6, p = 0.0075, FDR = 0.3980), DAPK1, IL24 and, IRF4 were downregulated in shAMBRA1 cells. Conversely, SEMA3C, JAG1, TCN1, WNT5A, FLT1, NRP1, TGFB1, TGFA, and TLR4 were upregulated (**Fig. 3B**). Genes involved in axon guidance, angiogenesis, cell-cell adhesion, cell migration and apoptosis were upregulated as a result of AMBRA1 knockdown (**Fig. 3C**). Western blot analysis confirmed the increased expression of FLT1 and WNT5A in A375 cells with AMBRA1 knockdown (**Fig. 3D, E**). STRING analysis showed that AMBRA1 loss leads to the upregulation of genes centered around MAPK activity (**Fig 4A**) and included TGFA, TGFB1, FGF2, FGF7, ITGA3 and NT5E. In addition, it upregulates FLT1, SEMA3A, SEMA3C, PGF and PDGFC which are key regulators of angiogenesis (**Fig 4B**). Collectively these data suggest the functional role of AMBRA1 extends beyond autophagy and cell proliferation regulation, and that AMBRA1 can regulate hallmark cellular pathways like angiogenesis and axon guidance.

**Fig. 4.**
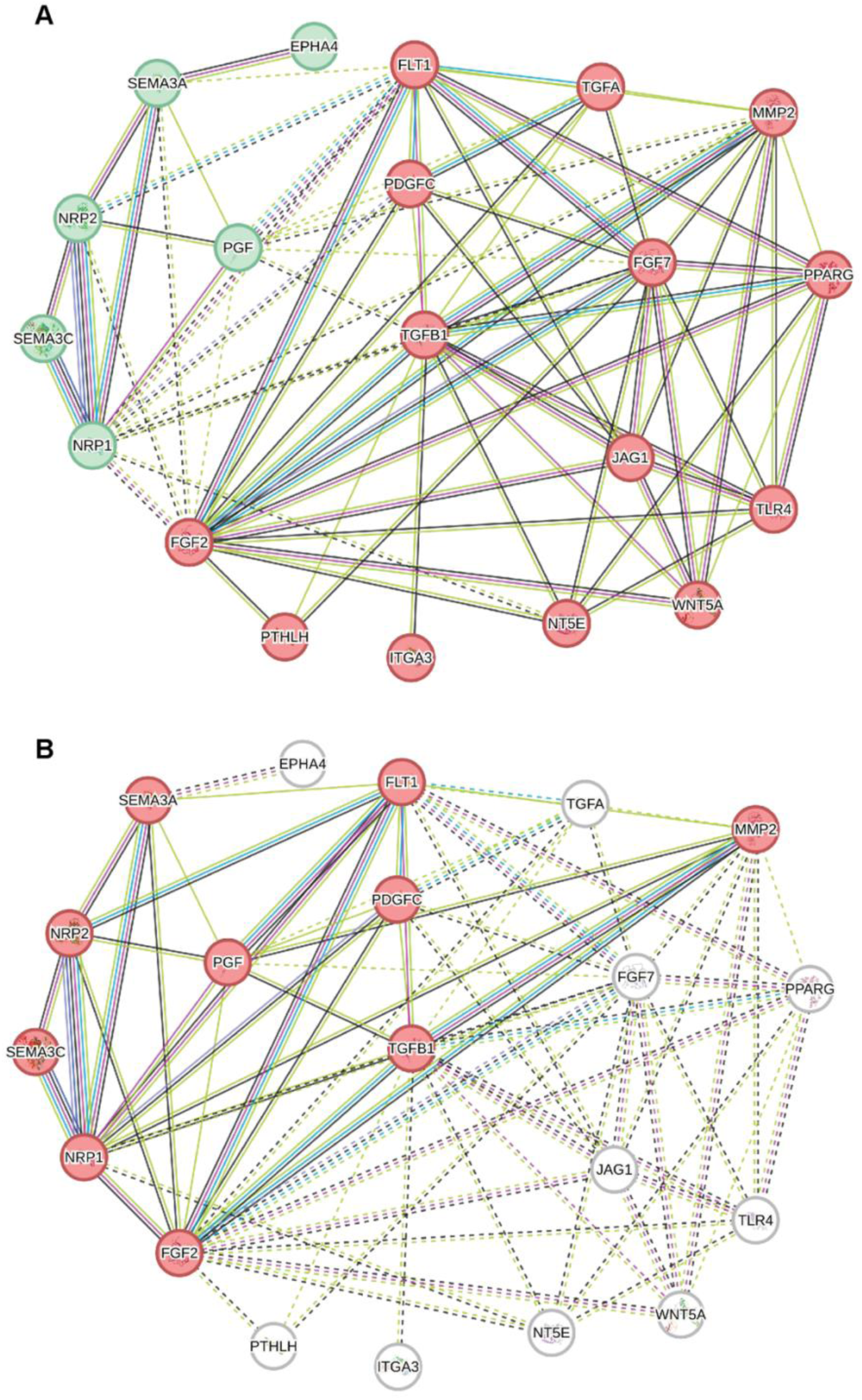
AMBRA1 loss leads to the activation of MAPK and angiogenesis signaling. (**A**) STRING protein network analysis showing shAMBRA1 overexpressed proteins network with those involved in MAPK signaling highlighted in red. (**B**) STRING protein network analysis showing shAMBRA1 overexpressed proteins network with those involved in angiogenesis signaling highlighted in red.

### Novel AMBRA1 binding partners

We next performed a yeast-two hybrid (Y2H) experiment using AMBRA1 as a bait to test for protein-protein interactions. We identified seven novel AMBRA1 protein binding partners: STX7, TMED7, ANK3, DSTN, AGO3 and ANKRD7. Of note, none of these interactions have been previously reported. We also used PPP2CA as a positive control as studies suggested that it can bind AMBRA1 [11]. Y2H experiments picked MAT2B, SHANK2 and PSMG2 as PPP2CA interactors but not AMBRA1, possibly due to limited number of screened positives. Collectively, we report novel AMBRA1 and PPP2CA binding partners that can constitute a larger network of proteins **(Fig. 5A**). To explore a more comprehensive AMBRA1 network, we searched reported AMBRA1 binding partners in the interactome BioPlex database (**Fig. 5B**) [18]. We then combined AMBRA1 binding partners identified in our study with partners previously reported in literature and in BioPlex. This integration resulted in a more complex protein network built around AMBRA1 (**Fig. 5C**), and this network confirms a role for AMBRA1 in autophagy and ubiquitination and suggests novel roles of AMBRA1 in the immune system and interleukin signaling (**Fig. 5D**). Of note, transcriptional analysis shows that AMBRA1 loss downregulates IL24 and IRF4 (**Fig. 3B**) further indicating a role of AMBRA1 in interleukin signaling and immune system.

**Fig. 5.**
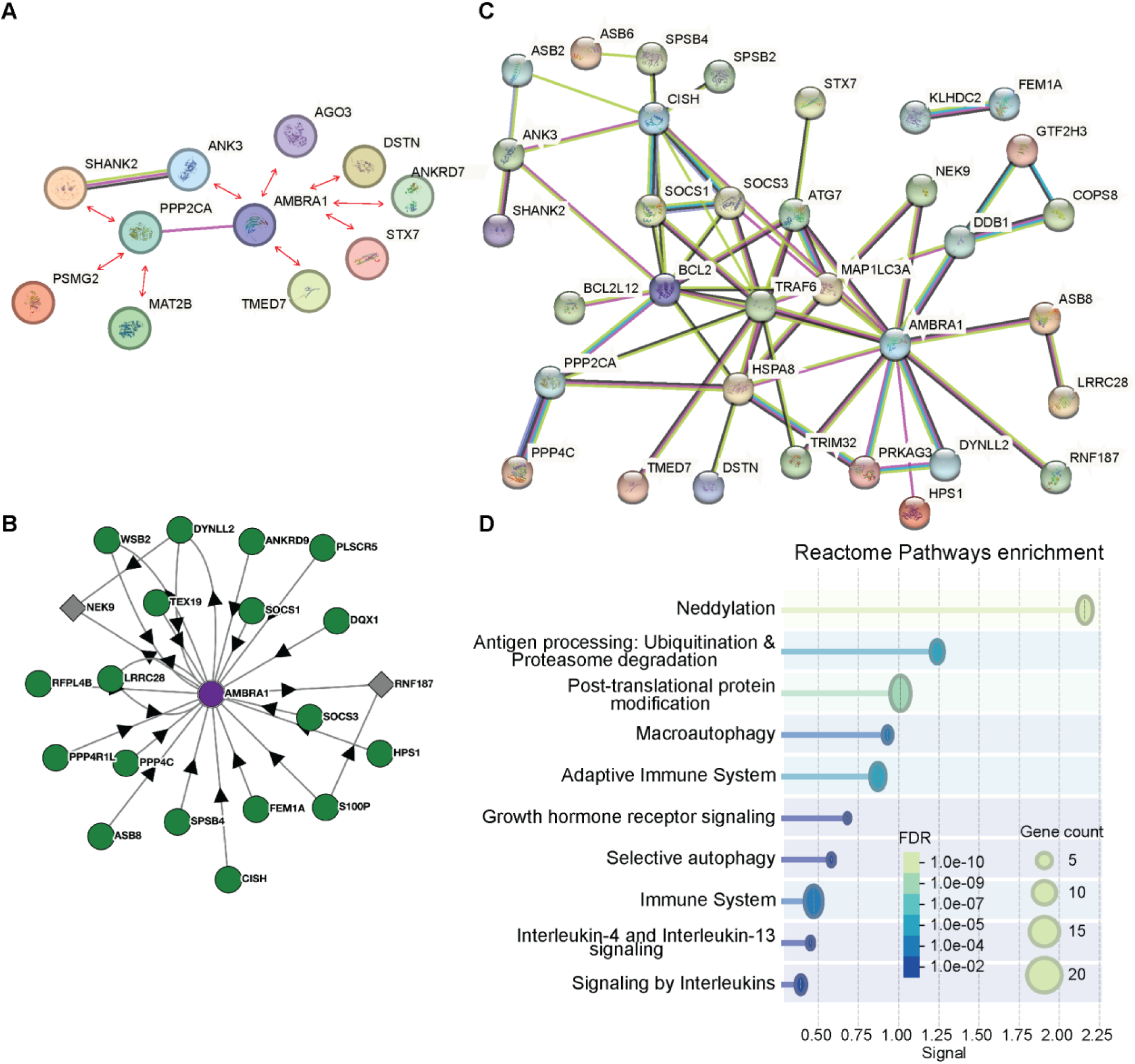
Novel AMBRA1 binding partners. (**A**) STRING protein interaction network of AMBRA1 and PPP2CA, novel interactions identified in this study are marked with red arrows. (**B**) Bioplex portal AMBRA1 binding partners. (**C**) STRING protein interaction network of AMBRA1 binding partners identified in literature, BioPlex and this study. Disconnected nodes were removed for simplifying. **(D)** Reactome pathway analysis of the protein network identified in (C).

## Discussion

Since its discovery in 2007, AMBRA1 research has significantly advanced with its reported role in regulating key biological pathways growing [6, 7, 10, 11]. Specifically, the role of *AMBRA1* in melanoma was highlighted in the recent years as it was shown that AMBRA1 loss in the epidermis surrounding melanoma can be a biomarker indicating worse prognosis [8, 9]. However, in other cancer types, the role of *AMBRA1* appears to be dynamic and can vary between one type of cancer and another [15, 17]. The mechanisms by which *AMBRA1* can play such roles remain largely unclear. Our study describes novel roles of *AMBRA1* that can potentially contribute, at least in part, to its role in melanoma and different cancer types.

Our bioinformatic analysis showed that AMBRA1 is lost in the skin adjacent to melanoma, a finding that was reported and confirmed in two recent studies [8, 9]. However, AMBRA1 levels were not different in metastatic compared to primary melanoma in four independent datasets. Moreover, TCGA data analysis suggest that *AMBRA1* mutations that lead to subsequent loss of expression are highly prevalent in melanoma. These data suggest that a subset of tumors could favor AMBRA1 loss. Our study suggests that tumors may favor AMBRA1 loss as it leads to the activation of MAPK signaling and angiogenesis related genes, providing the tumors with an advantage of increased proliferation and metastatic potential. Moreover, our results indicate that AMBRA1 loss leads to the overexpression of canonical Wnt signaling via FZD8 and non-canonical Wnt signaling via WNT5A and through binding ANK3, a regulator of Wnt signaling [19]. The Wnt family of proteins is involved in the regulation of cell proliferation, cell motility, cell polarity, organogenesis, cell fate and stem cells renewals [20]. WNT5A is a member of the Wnt family that signals through both the canonical and non-canonical Wnt pathways but, it is most often associated with non-canonical Wnt signaling [21]. The role of WNT5A in cancer remains under investigation [22] but evidence suggest that it can have a pro tumorigenic role in melanoma [23, 24]. In this study we report that in metastatic A375 melanoma cells the overexpression of AMBRA1 did not appear to influence the proliferation rate in an environment where nutrition is not limited. However, knockdown of AMBRA1 resulted in decreased cell proliferation. This result was not expected as AMBRA1 loss has been shown to promote melanoma growth and invasion [10]. However, analyzing Depmap portal showed that AMBRA1 gene knockdown or knockout can have dual effect on different melanoma cell lines, and it had a negative effect on the proliferation in A375 similar to what we observed in our study. A simple explanation is that the blockage of autophagy by AMBRA1 knockdown is causing a decreased proliferation as melanoma cells need the autophagy machinery to avoid apoptosis and cell death [25, 26]. We noticed that the cells favored AMBRA1 expression as cell proliferation was restored after prolonged time of shRNA selection removal.

The mechanism by which AMBRA1 can affect cell cycle and proliferation has been reported, at the post-translational level, to be through interaction with c-Myc and PPP2CA in a study that demonstrated elevated levels of Cyclins A and B upon AMBRA1 depletion [11]. Another study demonstrated that AMBRA1 can post translationally regulate Cyclin D degradation [12]. In this study that we have identified an interaction between AMBRA1 and ANKRD7, the latter is an effector of the small RAB GTPases RAB32 and RAB38 which can indicate a possible regulation of RAS pathway, a hall mark of cell proliferation [27], by AMBRA1. Also, AGO3 was among the proteins identified to interact with AMBRA1. This protein is reported to have a role in stem cell proliferation, regulating gene expression and it is essential in human embryogenesis [28]. AMBRA1 has also been reported to be implicated in embryogenesis [6]. Therefore, AMBRA1 and AGO3 might regulate one another during these key biological processes.

In addition, our study suggests novel roles of AMBRA1 in regulating multiple cellular pathways. For example, AMBRA1 was shown to be essential for nervous system development during embryogenesis and is highly expressed in various neural tissues [6]. Our study suggests a role for AMBRA1 in regulating axon guidance signaling, which is the process by which neuronal axons grow to reach their targets and establish connections, by altering the expression of key genes in this pathway. Knockdown of AMBRA1 significantly affects axon guidance. Additionally, ANK3, an identified AMBBRA1 binding partner, is involved in axon segment initiation. Moreover, our study reports that AMBRA1 can be central in a larger protein network that can affect immune response, suggesting that knocking down AMBRA1 in vitro can limit the scope of studying its effect on cell proliferation.

In summary, our study reveals potential roles of AMBRA1 that can extend beyond autophagy and its role reported in cell proliferation and it identifies protein interactors of AMBRA1 that can regulate or be regulated through this interaction. Our study suggests that AMBRA1 loss in some melanoma tumors can result in the upregulation of angiogenic and metastatic genes that can attribute to its role as tumor suppressor in melanoma.

## Materials and Methods

### Bioinformatic analysis

Normalized counts were downloaded from GEO datasets (GSE15605 and GSE59499) or TCGA SKCM. Gepia2 [29] was used for survival analysis and cbioportal [30] was used for mutation analysis.

### Cell cultures

Cells were cultured and routinely passaged to be maintained in the exponential phase in either high or low Dulbecco’s Modified Eagle Medium (DMEM) (Bio-sera). The media was supplemented with 10% Foetal Bovine serum (Gibco), 100 µg/ml primocin (Invivogen) and 300 µg/ml L-glutamine (Lab-tech). Cells were grown at 37°C in a 5% CO2 humidified atmosphere.

Transfection maintenance was performed by adding antibiotics to the media. The overexpression strains were maintained using G418 at a concentration of 2 µg ml-1 and the knock-down cell lines were maintained using puromycin at a concentration of 2 µg ml-1.

### Lentiviral production and infection

Lentiviral particles were generated by co-transfecting 293T packaging cells with a total of 10 μg of shRNA-expressing pLKO vectors targeting *AMBRA1* (three different constructs, each with a unique targeting sequence; Merck), 2.5 μg of a vesicular stomatitis virus G (VSV-G) envelope expression plasmid, and the psPAX2 packaging plasmid (which encodes the *gag*, *pol*, and *env* genes). Transfections were carried out using the calcium phosphate precipitation method [31]. At 48 hours post-transfection, the culture supernatants containing lentiviral particles were harvested and filtered through a 0.45-μm pore filter to remove cellular debris. To enhance transduction efficiency, Polybrene was added to a final concentration of 4 μg/mL. The viral supernatant was then used to infect A375 cells. Following selection, the knockdown efficiency of each shRNA construct was evaluated, and the most effective shRNA(s) were selected for use in subsequent experimental assays.

### Retroviral production and infection

Retroviral particles were produced by co-transfecting 293 gp/bsr packaging cells with 15 μg of retroviral vectors encoding AMBRA1 (overexpression construct) [32] and 5 μg of a vesicular stomatitis virus G (VSV-G) envelope protein expression plasmid. Transfection was carried out using the calcium phosphate precipitation method. Forty-eight hours post-transfection, culture supernatants containing retroviral particles were collected, filtered, and supplemented with 4 μg/mL polybrene to enhance infection efficiency. A375 cells were then transduced by incubating them with the viral supernatant for 6–8 hours. Overexpression of AMBRA1 in infected A375 cells was subsequently confirmed by quantitative PCR (qPCR).

### Incucyte live cell growth assay

2000 cells/well were seeded in 96 well plates. Nine wells were used as replicates for each STCs and/or treatment or condition. Live cells were imaged using the IncuCyte system (Essen BioScience), taking four images per well every 2 hours for 5 days after the cells were seeded. IncuCyte software was used to segment and process object count (cells per image) and represent it as the average number of cells per image over time. Data are presented as mean+/− standard deviation. p-values were calculated using an independent, two-sided t-test by comparing the non-linear fit of exponential growth of each cell line against its control, and significance was calculated as p<0.05

### Western blot analysis

Cells were seeded at a concentration of 5×10^5^ to a 6-well plate and grown at 37°C and maintained in the exponential phase. Cells were then washed twice with PBS and harvested with 200 µl 1x cell lysis buffer (Abcam) and sonicated for 3×10 seconds. Lysates were stored at –80°C and used for downstream analysis. Protein quantification was performed using BSA as a standard and Bradford reagent according to the manufacturer’s instructions. Approximately 25 µg of protein was resolved on a 12% Stain free TGX polyacrylamide gel (Bio-Rad) electrophoresis. A stain free fluorescent image of the gel was captured before gels were transferred to polyvinylidene difluoride (PVDF) using the transblot turbo system (Bio-Rad). Membranes were blocked in Immobilon® Block-Chemiluminescent Blocker (CB) (Merck) for 1 hour at room temperature. Membranes were then incubated with primary antibodies overnight at 4°C with shaking, washed 3×10 minutes with TBST and incubated with secondary antibodies for 1 hour at room temperature followed by another 3×10 minutes wash, membranes were then incubated with 2 ml freshly prepared (1:1) mixture of peroxide reagent and luminol/enhancer reagent supplied from Bio-Rad for chemiluminescent detection of HRP activity of the conjugated secondary antibody. The membranes were then analyzed by the ChemiDoc™ Imaging System (Bio-Rad, UK) for protein band detection. Antibodies used: Rabbit anti human AMBRA1 antibody (Covalab 00013214), Chicken anti-beta-Galactosidase (Sigma GW20071F), Mouse monoclonal to FLT1 (VEGF receptor 1) (Abcam, ab9540), Rabbit monoclonal to Wnt5a-C-terminal (Abcam, ab235966), Goat anti-chicken IGY (Abcam, ab97135), Goat anti rabbit IgG (Vector labs, AI-1000-1.5) and Goat anti mouse IgG (Vector labs, AI-9200-1.5). Quantitative analysis was performed using Imagelab version 4.0 (Bio-Rad). Pixel intensities of western bands were quantified by creating identical volume boxes and using a separate background volume to perform background subtraction. Total protein loading per lane was quantified by generating the pixel intensities of the protein loading from the stain free gel image as outlined above. The relative pixel intensities per lane were normalized to a nominal total protein lane set as a nominal value of 1. The normalized relative pixel intensities of the western bands were calculated as their band intensities divided by the normalized protein loading value. The use of stain free gels for protein normalization has been shown to have advantages over performing a separate loading control [33, 34].

### RNA extraction and quality analysis

Total RNA was extracted from each cell line using Trizol reagent (Thermo-Fisher, Cat. 15596026) according to the manufacturer’s instructions and the precipitated RNA was re-suspended in 200 µl 1xRNA secure reagent (Ambion). RNA quality analysis was performed using Experion™ RNA StdSens and HighSens Analysis Kits from Bio-Rad following instructions from the manufacturer.

### Gene expression analysis by microarray

The analysis was performed by Source bio-Science. Three biological replicates from each cell line at a concentration of 120 ng/µl were hybridized on GeneChip™ Human Gene 2.0 ST.The microarray data was normalized and analyzed using Transcriptome analysis suite (TAC version 4.0.1) (Thermo-fisher). Ebayes Anova method was used, and the analysis was set to a P-Value < 0.05. Functional analyses were performed uing Gene Ontology (GO) resource [35, 36] and STRING protein-protein interaction network [37, 38]

### Yeast-Two Hybrid Protein-Protein interaction

Yeast strains were maintained by streaking at least once every 4 weeks on fresh YPDA plates, incubated at 30°C for 3 to 5 days then stored at 4°C. Only fresh grown colonies were used for transformations. Plates were prepared by adding each media pouch content to 500 ml deionized water then autoclaving, left to cool to 50°C then X-α-gal and/or antibiotics were added before pouring the plates. Yeast transformation, positive and negative control experiments were performed as stated by Yeast-Two Hybrid System manufacturer (Clontech Laboratories). All isolated plasmids were sequenced using a t7 promoter at Source Bio-science limited.

Yeast transformation, positive and negative control experiments were performed as stated by Yeast-Two Hybrid System manufacturer (Clontech Laboratories) a positive control mating was performed by mating Y2H gold strain fused with murine p53 and Y187 yeast strain fused with SV40 large T-antigen. On the other hand, a negative control experiment was performed by mating Y2h gold yeast stain fused with lamin and Y187 yeast strain fused with SV40 large T-antigen.

### cDNA synthesis and PCR amplification

PPP2CA and AMBRA1 cDNAs were prepared using oligo primers by PCR using RNA extracted from U-937 (ATCC® CRL-1593.2™) and A-375 (ATCC® CRL-1619™) cell lines. An ORF clone with AMBRA1 sequence was purchased to be used as a PCR template. PCR reactions were performed on Bio-Rad T100 thermal cycle. PCR reactions were performed using 10 x Immomix master mixes (Bio-Rad), Q5 polymerase (Bio-labs), and proof-reading polymerase was later used (Iproof Bio-Rad).

Primers:

PPP2CA-Forward 5’-CGC GAA TTC ATG GAC GAG AAG GTG TTC ACC –3’ PPP2CA-Reverse 5’-CGC GGA TCC TTA CAG GAA GTA GTC TGG GGT ACG –3’ AMBRA1-Forward 5’-AGG AGG ACC TGC ATA TGA AGG TTG TCC CAG AAA AGA ATG CC –3’ AMBRA1-Reverse 5’ GCC TCC ATG GCC ATA CTA CCT GTT CCG TGG TTC TCC C-3’

### DNA digestion, Ligation, Plasmid extraction from E. coli and transformation into yeast cells

Plasmids were transformed to competent E. coli (DH5-α strain, Invitrogen 18258012) according to manufacturer’s protocol using appropriate antibiotic for selection. Plasmids were transformed into the yeast strains according to the manufacture specifications (Clontech). Plasmids were extracted from E. coli using plasmid plus midi kit (QIAGEN technologies) according to manufacturer’s instructions. All of the PCR products and plasmids were digested by using 200 ng DNA to 1 µl of the restriction enzymes (*BamH* I, *EcoR* I, or *Nde* I, Thermo Scientifc). All ligation reactions were made at a vector to insert ratio of 1:5, the appropriate ligase was added to the reaction mixture and incubated overnight at room temperature.

## Statistical analyses

Statistical analyses were performed with Microsoft Excel, GraphPad Prism (version 10.2.0, RRID:SCR_002798), or R statistical software (RRID:SCR_001905 version 4.3.1).

## Data Availability

Microarray transcriptomic data are available at GEO datasets (GSE175361)

