## Supplementary material for "Investigating the relationship Between AMBRA1 and Cell Proliferation in Melanoma": Figure S1

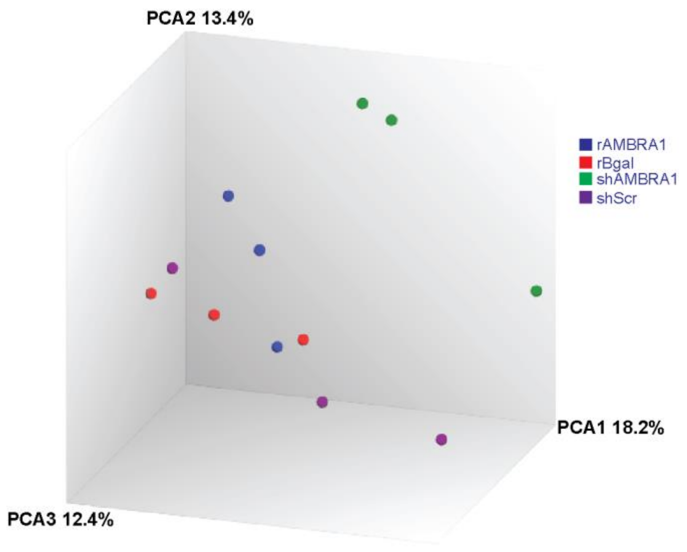


**Fig. S1 AMBRA1 regulate hallmark cellular processes on the transcriptional level**

(**A**) PCA plot of cells overexpressing AMBRA1 (rAMBRA), overexpression control (rBgal), AMBRA1 knockdown (shAMBRA1), and knockdown control (shScramble).
